# A longer stance is more stable for a standing horse

**DOI:** 10.1101/2021.10.15.464610

**Authors:** Karen Gellman, Andy Ruina

**Affiliations:** Maximum HorsePower Research, Nottingham, PA 19362; Mechanical Engineering, Cornell University, Ithaca, NY 14853

**Keywords:** statics, horse, posture, balance, equilibrium, stability, mechanism

## Abstract

What is the effect of posture on the stability of a standing horse? We address this with a 2D quasistatic model. The model horse has 3 rigid parts: a trunk, a massless fore-limb and a massless rear limb, and has hinges at the shoulder, hip, and hooves. The postural parameter *ℓ_g_* is the distance between the hooves. For a given *ℓ_g_*, statics finds an equilibrium configuration which, with no muscle stabilization, is unstable. To measure the neuro-muscular effort to maintain stability, we add springs at the shoulder and hip; the larger the springs needed to stabilize the model, the more the neuro-muscular effort needed for stabilization. We find that a canted-in posture (small *ℓ_g_*), observed in some pathological domestic horses, requires about twice the spring stiffness (representing twice the neuromuscular effort) as is needed for postures with vertical or slightly splayed-out (large *ℓ_g_*) legs.

## 1 Introduction

The assumption that horses tend to minimize metabolic cost predicts some equine locomotion features (*e.g.*, gait transitions) [13, 10, 17, 14]. However, horses typically spending much more time (up to 23 hours a day), standing than on the move. They eat, socialize and sleep while standing. While the metabolic costs associated with locomotion are obvious, it also takes some effort to stand still. This paper uses a simple model to explore the idea that, as for locomotion, horses choose ways of standing that are relatively easy for them.

There are few studies of standing horse posture. Horse postural sway has been characterized with stabilograms (trajectories of the net center of pressure) [5, 6]. As with humans [16], the clinical utility of using stabilograms to measure postural competence is likely limited. Lesimple, Fureix et al [9, 18] found that a horse with high head height, relative to back height, tended to be in stress or pain. Physiotherapists claim a correlation between equine thoracolumbar spinal contours and back pain [26, 28]. Textbooks (*e.g.*, [2]) (incorrectly, we believe) describe some postures as ‘conformations’, categorizing these postures as built-in physical features of the horses body rather than as outputs of neuromuscular control.

Feral horses living in a natural unconfined environment, and most appropriately-managed domestic horses, are usually seen with visually-vertical limbs. There are various reasons that long-legged animals might choose to stand with vertical, as opposed to canted-in or splayed-out, legs. (See Fig.1)

**Figure 1:**
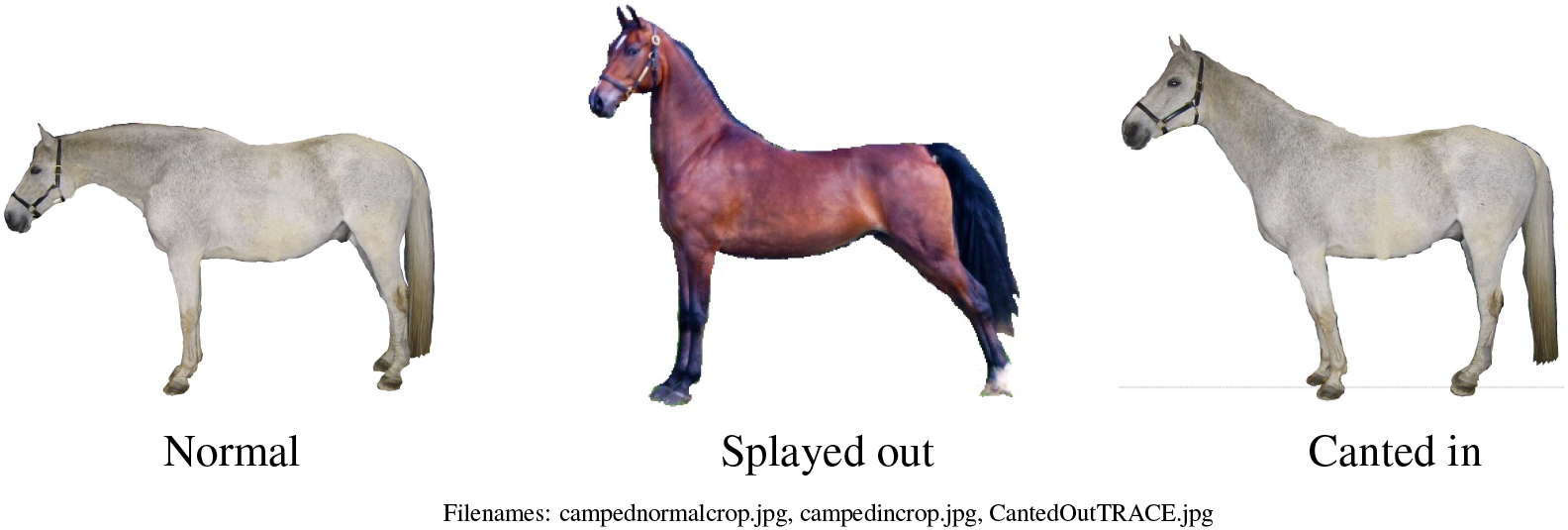
Horse postures. **Left:** *Normal*, neutral horse posture, legs are approximately vertical; **Center:** *Splayed-out;* the the fore and hind feet are relatively spread; and **Right:** *Canted-in*, the fore hoofs and hind hoofs are closer together.

1. The compression force in the legs is minimized by having vertical, as opposed to sloped, legs (see Methods);
2. In locomotion, the range of leg motion is roughly centered on vertical (protraction and retraction of the limb are approximately symmetric relative to vertical). So, neither agonist nor antagonist muscles at the shoulder or hip are near their maximum (possibly uncomfortable) lengths when the leg is vertical.
3. The peak vertical ground reaction forces in running gaits occur when the legs are near to vertical, so the bones, muscles, tendons and ligaments are configured for carrying loads with vertical legs. Conversely, standing with vertical legs might help align the internal structures for the running leg loads;
4. Horses are prey animals; a horse with vertical legs is perhaps most ready to step in any direction.

Yet, horses do sometimes choose splayed-out or canted-in postures.

Horses choose a splayed-out posture when carrying a heavy rider, when holding a heavy cart stationary (’parked’), when a mare is in the late stages of pregnancy, or in the first hours that a newborn foal stands. The intuitive idea, which we buttress with our model here, is that a splayed-out posture is more stable.

On the other hand, canted-in postures, in which horses stand with their legs relatively close together beneath their trunk, are seen in some horses with chronic performance deficits (J.M. Shoemaker, DVM — personal communication), in horses with Equine Motor Neuron Disease (EMND, [7, 29]), and, perhaps, in extreme fatigue (Appendix G). In contrast to the possible stability gain from a splayed-out postures, the possible reasons some horses choose a canted-in posture are less obvious. Perhaps,

1. The horse has distorted proprioception so misperceives its body position, and holds inappropriate posture, which happens to be canted-in;
2. The horse has an impairment of postural control or motor output such that it is unable to stand with vertical or splayed out legs;
3. The canted in posture allows the horse to ‘lean’ on ligaments or bones in ways that may be painful or injurious, but relieve the burden on impaired muscles (see Appendix G);
4. Following the model of Scrivens *et al* [25] for human standing, a canted-in posture may be easier to stabilize if selecting control gains is the primary neuro-muscular challenge and neural delays are non-neglgible (see Appendix E).

Our approach to understanding postural choice here, is to study the statics and stability of standing postures. In particular, how does a horse’s choice of leg splay (vertical vs splayed-out vs canted-in) affect the difficulty of maintaining stability?

Our approach is the same as some have used for stability of grasp (*e.g.*, [27, 21]), except that we have a more general interpretation of stiffness which includes stiffness that is approximated by quick linear feedback. Our approach is a simpler statics calculation for the same model pursued by [25, 3] and [12] using dynamics; their work also incorporates neural time delays, which we neglect (see Methods, Discussion and Appendix E).

As noted in [12], the simple idea that a broader stance is more stable because the base of support (the polygon defined by the hoof positions) is bigger, is not key. In our model, an uncontrolled standing horse is a unstable mechanism, even when there is no danger of tipping over like a plastic horse.

There is no single obvious way to quantify the “difficulty of maintaining balance.” The appropriate measure of difficulty depends on the nature of the employed control system. In particular, the torques used to stabilize an equilibrium posture may come from tonic contractions or co-contractions (e.g., [22]), from proportional feedback with delays (e.g., [12]), or from intermittent feedback based on an internal model [20]. Here, we assume that the tonic component is either significant in itself, or is a reasonable proxy (for our purposes) of the other mechanisms. We quantify degree of instability by the minimum stiffness *k*_min_ of shoulder and hip springs that, if added, could stabilize an equilibrium posture. Use of *k*_min_ as a proxy for neuro-muscular control effort may be regarded narrowly as measuring the difficulty of maintaining balance with tonic contractions alone. Or, *k*_min_ might be interpreted more broadly as representative of the challenge of control by all mechanisms combined. Our use of *k*_min_ as a proxy for neuro-muscular control difficulty would be inappropriate if neural delays are actually the primary control challenge for standing horses. These issues are further discussed in Appendix E. The minimum stiffness *k*_min_ is equivalent to the minimum needed gain, if both the sensing and torque output are at the shoulder and hip joints.

## 2 Methods

We make the common biomechanical approximation that a whole animal is a linkage of rigid parts connected by hinges. During quiet stance, the lower leg-joints of the weight-bearing legs are generally locked in an extended (generally straight) configuration, and there is little deformation of the back. So, we assume that stabilizing the spine, and holding the legs straight, is either non-problematic for a horse, or is an equal problem independent of stance width. Hence, the deformations of interest here are only the rotations at the shoulder and hip joints. By “shoulder” and “hip” we henceforth mean the effective centers of rotation of the model legs relative to the body (See Appendix F for specific parameters of the model). Standing involves relatively slow motions, so our model uses statics. See Appendix A for discussion of the neglect of ‘inertial forces’ (note, again, our simplification of dynamics to statics is in contrast to the dynamics postural models of [25, 3] and [12]). Because the legs are much lighter than the body, for simplicity, we neglect the weight of the legs. Thus, we consider the statics of a two-dimensional horse consisting of a rigid body (trunk, neck and head), with mass, that is supported by negligible-mass, rigid fore-legs and hind legs.

For simplicity, and because it is most relevant to our observations of horses, we consider the sagittal plane (side view) only. The sagittal plane horse model here is similar to the frontal plane models used for analysis of standing cats and humans used by [25, 3] and [12]. As in [25, 3, 12], the model hwew is a ‘four-bar’ linkage. Like [25, 3], our model has no other moving parts (in contrast, [12] uses a similar model that also has an independently moving upper body). The frontal-plane models of [3] and [12] are left-right symmetrical. In contrast, our horse model does not have that symmetry (front is different from back). Using this 3-link model (4-bar linkage, including the ground as a link), we consider two statics features:

### 1) Geometry of Equilibrium

For a given foot spacing, basic statics (Appendix B) finds those postures for which the linkage carries the whole load without joint torques (assuming a perfect model and no disturbances). In these joint-torque-free postures the ‘horse’ (meaning our model of a horse) and all of its parts obey the laws of static equilibrium (*i.e.*, forces and moments ‘balance’ on all of the parts). For symmetric models, as in [3] and [12], this posture is found by inspection as the symmetric posture. Because a horse is not front-back symmetric, nor is our model, finding equilibrium postures requires a statics calculation.

### 2) Stability of equilibrium

An equilibrium posture, as found above, can be stable or unstable (Appendix C). An unstable equilibrium posture will, if disturbed slightly, tend to deviate progressively. In contrast, a stable equilibrium posture is one that, if slightly disturbed, will spontaneously tend to recover. Here, we want to quantify the degree of instability of an unstable posture. We seek a measure of the neuro-muscular effort needed for stabilization. Our proxy for this is the amount of joint torque needed to bring the horse back to equilibrium after a given small disturbance. In particular we quantify instability by the stiffness of the springs at the shoulder and hip needed to achieve stability. As noted above, we differ from [25, 3] in our quantification of this effort (Appendix E).

The distinction between an equilibrium posture and a stable equilibrium posture, can be understood by analogy with a pendulum (a stick hinged at one end). A pendulum is in equilibrium if pointed straight down or if pointed straight up. However, only straight down is a stable equilibrium. The unstable, straight-up equilibrium of a pendulum can be stabilized by adding correction (a force or torque that rights the pendulum when it deviates from upright). For example, a sufficiently stiff torsional spring at the hinge will stabilize the upright, otherwise unstable, equilibrium. Since a longer or heavier upright pendulum needs a stiffer spring to be stabilized, it might be regarded as more unstable than a shorter, or lighter pendulum, if we measure instability by the stiffness of the minimal stabilizing spring (see Appendix D for expansion of this pendulum analogy and implications for various candidate quantifications of instability).

The equilibrium postures are found using the statics (force and moment balance) of each leg, and the statics of the whole ‘horse’, as described in detail in Appendix B. Stability is determined by whether the equilibrium is, or is not, at a potential energy minimum. The gravitational potential energy is calculated numerically for postures that are close to the equilibrium postures. For unstable postures, the degree of instability is quantified, in our model, by stabilizing the horse with equal springs at the shoulder and hip. As noted, these springs are proxy for neuromuscular effort in the living horse. The spring potential-energy is then added to the gravitational potential energy. We find the smallest springs (*k*_min_) necessary to bring the net potential energy to a local minimum. We then assess degree of stability by looking at the needed stiffness to stabilize a horse with a given leg splay. The details of these calculations are described in Appendix C.

### Leg compression force

In this analysis, we lose little accuracy by replacing our model with one in which the fore and hind legs have equal length. The compression in the legs, *F_c_*, is then proportional to *W/* cos *θ*, where *W* is the weight born by the leg and *θ* is the deviation from vertical. For small angles, this is approximately quadratic in leg angle, *F_c_* ≈ *W*(1 + *θ*^2^/2) (where *θ* is measured in radians), and is not a dramatic effect. For example, for *θ* = 15° the compression is increased by 3.5% compared to having vertical legs (*θ* = 0°). Canted-in or splayed-out limb angles seen in living horses are usually under *θ* = 15° [11].

## 3 Results

### The muscle-free equilibrium postures

have a simple geometric interpretation (Fig. 2). Consider a vertical line through the center of mass; a line defined by the front hooves and shoulder; and a line defined by the rear hooves and hip. Because the legs are ‘two-force-objects’ and the body is a ‘three-force object’ these three lines are either parallel (Fig. 2a, and Appendix B), or they intersect above or below the horse at C (Figs. 2b and c). These postures, if exact, don’t need joint torques to be maintained. Thus, we expect the postures that horses adopt would be close to these. For example, if the legs are parallel, they should be vertical (not slanted) with respect to the ground — making a rectangle. A horse with parallel slanted legs is not in a muscle-free equilibrium posture.

**Figure 2:**
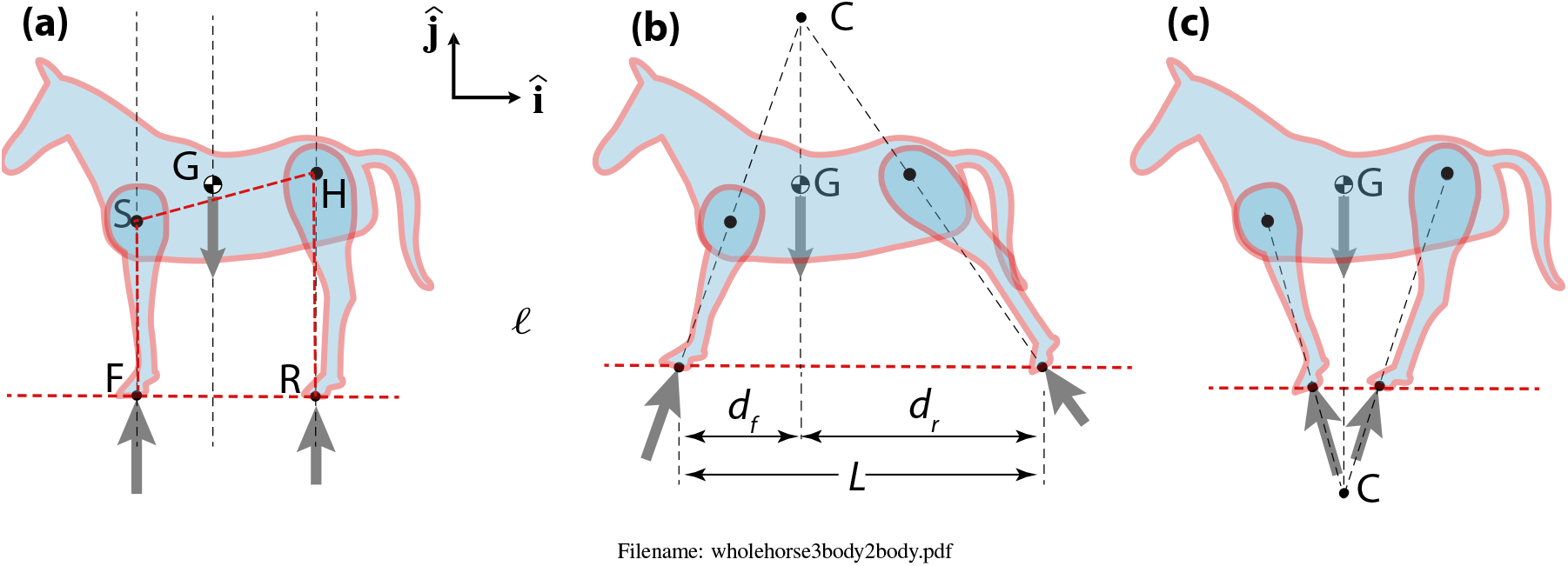
Three muscle-free equilibrium postures: **(a)** Vertical legs; **(b)** Splayed-out; and **(c)** Canted-in. In our 2D model, a whole horse has three forces acing on it: 1) gravity; 2) the ground reaction force on the hind feet; and 3) the ground reaction force on the fore feet. Static equilibrium of the whole horse, a ‘three-force object’, demands that these ground forces are either **(a)** parallel and vertical; or **(b)** they intersect directly above the center of mass (CoM); or **(c)** they intersect directly below the CoM (see appendix B for reasoning). The corrective joint torques needed to stabilize these equilibria are considerred in Fig. 3.

### Degree of stability

For an standing horse, physically-realizable muscle-free equilibrium postures are all unstable; that is, the equilibrium postures are postures in which small deviations from equilibrium would tend to grow rather than be naturally restored. In other words, for all of the equilibrium postures of interest, the center of mass height is at a (local) maximum. This instability occurs whether the leg-lines intersection point C is above or below the horse. Here, we add springs at teh hip and shoulder and find *k*_min_, the smallest spring that stabilizes the muscle-free equilibrium postures. Our basic modeling postulate is this:

> *A posture with a bigger k*_min_ *is more unstable*.

The central result of the calculations, found in the appendices, are shown in Fig. 3:

> *A horse, when legs are canted in (legs closer together) needs much larger corrective springs (by more than a factor of 2) than when legs are vertical or slightly splayed out (normal neutral posture, NNP).*

**Figure 3:**
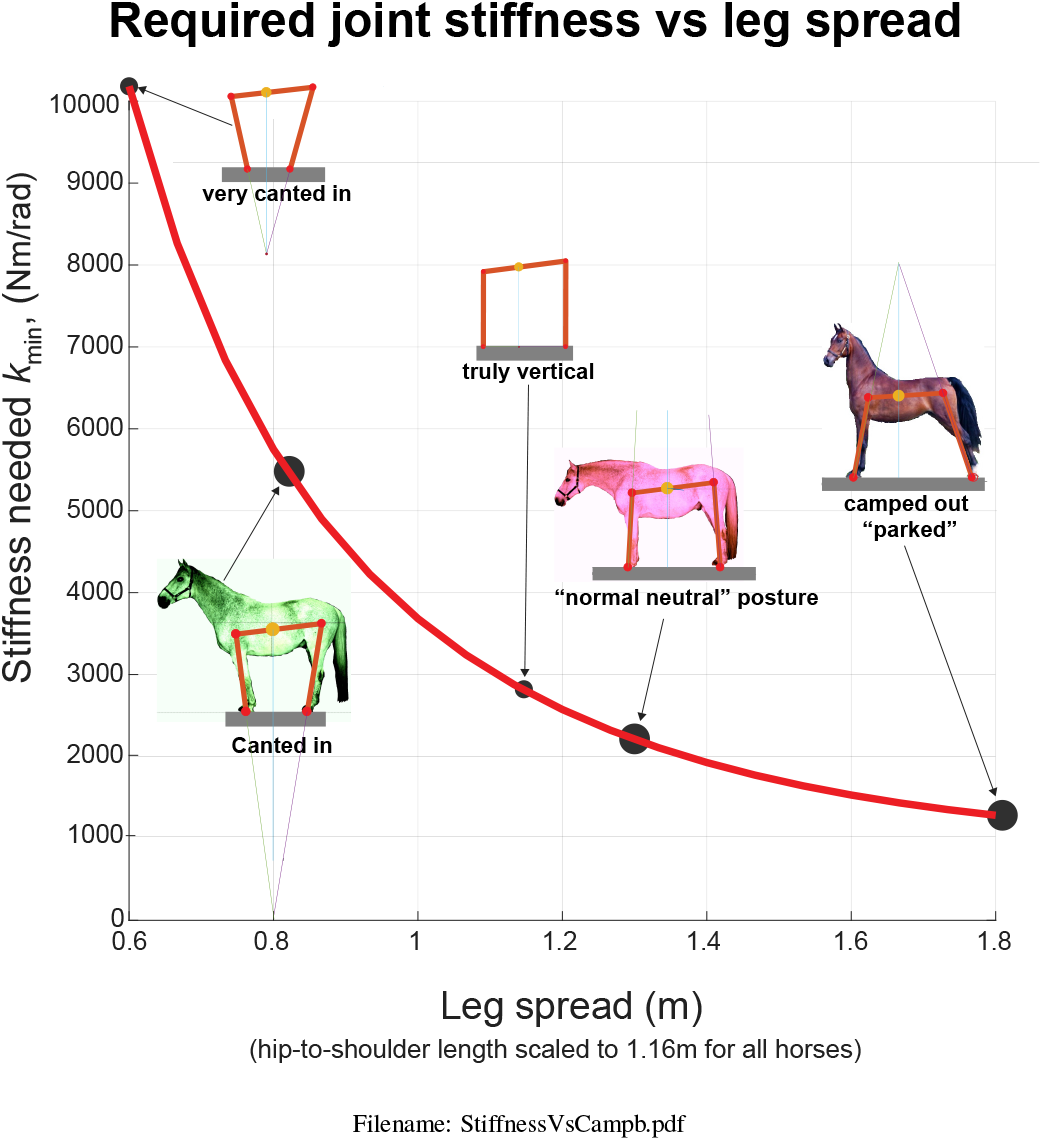
Stiffness vs Leg Splay. For a given horse geometry (leg lengths, back length, location of CoM) and for a given splay (distance between fore and hind hooves) the ‘horse’ (modeled as a linkage) has an equilibrium configuration satisfying the statics three-force-object rules for the horse and the statics two-force-object rules for the legs (see Fig. 2). This equilibrium can be stabilized with equal springs at the shoulder and hip. The size *k*_min_ of the minimum spring is plotted as a function of leg splay. The central results of this paper are the differences between *k*_min_ for the normal neutral posture and for the splayed-out and canted-in postures. To be stable, the canted-in posture needs a (spring) stiffness (*k*_min_ ≈ 5400 Nm/rad) that is more than twice that needed for the normal neutral posture (*k*_min_ ≈ 2100 Nm/rad for NNP). The splayed out posture is even more intrinsically stable than NNP in that it needs only about 60% of the *k*_min_ required by NNP.

Although the 4-bar linkage model of a human in [3] differs in quantitative detail (relatively shorter body, reflection symmetry) from our horse model, the mechanics seems to show all the same trends as our model. In particular, [3] noted, as is true here, that the joints have more mechanical advantage for supporting the center of mass in the splayed-out configuration (see appendix C).

A final statics result is that Within the commonly observed range of horse postures, the compressive forces in the legs, while minimized by vertical legs, are only slightly increased with canted-in or splayed-out legs.

## 4 Discussion

In normal, sound horses the most common posture is standing with metacarpal and metatarsal bones visually vertical. In this posture the legs are slightly splayed-out with reference to the shoulder/hip to hoof angle, as seen in Fig. 3. We term this normal neutral posture (NNP): <u>n</u>ormal in that it is most common in healthy and feral horses; <u>n</u>eutral in that it equally allows leg movement in any direction, <u>p</u>osture. Our model predicts greater stability in splayed-out postures, thus possibly explaining

- Why horses needing an extra degree of stability often display splayed-out postures; and
- Why we rarely observe persistent canted-in postures in normal healthy situations.

Like horses, humans adopt a wider stance when lifting heavy weights or standing on unstable surfaces.

Why don’t humans and quadrupeds always use a more-spread stance? Stability is only one of several situationally-dependent goals. In Fig. 3, the leg compression changes minimally between NNP (slightly splayed) and more obviously splayed-out, and more stable, posture. Perhaps the various benefits of nearly vertical legs outweight the stability-benefits of a wider splayed-out stance.

Why, then, are canted-in postures ever seen in horses (or people)? Narrow-stance postures seem to be associated with neural impairment with sensory input, neural processing or motor output concerning postural control (See Appendix G for speculation on another cause of collapsed posture).

1. Parkinson’s Disease (PD) patients (humans) have a characteristic collapsed standing posture that includes having a narrow stance. Parkinsons is a degenerative disorder of the human central nervous system [15] in which patients have lost dopaminergic neurons in the midbrain, a critical region for postural control, resulting in a dysfunction of central postural processing.
2. Equine Motor Neuron Disease (EMND) is a condition for which canted-in limb posture is pathognomonic (characteristic) in horses. In this degenerative neuropathy, similar to ALS in humans, there is cell death and consequent denervation atrophy in postural muscles [29]. EMND patients habitually display extreme canted-in postures, fatigue quickly and spend much horse time lying down [7]. This is a defect in the postural motor output.
3. Abnormal Compensatory Posture (ACP), a narrow stance seen in some domesticated horses, appears to be caused by structural distortions in some critical mechanoreceptor-rich anatomic regions (the hooves, the stomatognathic system, and the upper cervical area) due to practices associated with domestication. In these horses, the inaccurate proprioceptive signals are altering neural processing and motor output. Horses habitually standing with canted-in posture often develop a predictable set of musculoskeletal problems, including: navicular syndrome, negative palmar/plantar angles, digital cushion degeneration, hock and stifle osteoarthritis, sacroiliac fixation or hypermobility, back pain, “kissing spines”, neck pain, and more (Judith M. Shoemaker DVM — personal communication). Their posture reverts to a visually vertical stance (NNP) when these structural and functional distortions are corrected [11]. ACP is a defect in sensory (proprioceptive) input.

As noted, Bingham *et al* [3] use a mechanical model, similar to ours but come to the opposite conclusion. They predict that a narrow-stance (canted-in) posture is ‘more stable’ whereas we find it is less stable. Bingham et al [3] argue that narrow stance might benefit human PD patients because, in their model the range of feedback gains which stabilize a narrow stance is greater than the range of gains that can stabilize a wider stance. Thus, they claim, finding a successful stabilizing gain is easier for a narrow rather than for a wider stance (see Appendix E for further comparisons with [3]).

Neither zoologists nor veterinarians have an accepted standard for ‘normal’ standing horse posture. Because standing is so much of a horses life, perhaps evaluation of habitual posture should be included as part of any clinical exam, or in developing a management program. The slightly-splayed-out NNP posture perhaps represents a trade-off between the stability benefits of more extreme splay and *e.g.*, the quick-escape benefits of mostly vertical legs. On the other hand, sports injury, poor athletic performance and chronic or recurring lameness are common sequelae in domestic horses who habitually display an abnormal compensatory posture (ACP) of canted-in limbs rather than a normal neutral posture (NNP). Future studies will aim to characterize the anatomic distortions associated with ACP and track restoration of NNP through correcting these structures.

## Acknowledgments

We thank Judith M. Shoemaker, DVM for clinical commentary on performance horses and editorial input, Elizabeth Reese, CTAT, MEd for discussions on posture and movement in humans, and Carl DeStefano, DC, DACNB, FACFN for discussions on CNS patterning. Robert Paterka, Lena Ting, Art Kuo, and Madhusudhan Venkadesan, all knowledgeable of the dynamics of standing still, made comments that helped us place our simple model in context.

## Author contributions

KG posed the question and provided concepts of posture for horses. AR made the figures, the numerical model and wrote the appendices. KG and AR contributed equally to the main text.

## Supplemental Appendices

These appendices are not intended to be in the journal, but are for an electronic supplement. These have

- Details that need mention, but which would distract from the main text;
- Expanded discussions of some topics; and some
- Pedagogical explanations.

## Appendix A: Basic modeling issues

In this appendix we review some of the reasoning behind, and concepts related to, the basic model features we use.

### A horse is a linkage

The common biomechanical model of a large terrestrial vertebrate is a linkage of rigid parts. For example, each leg is modeled as a set of rigid links, with each link representing a bone and the flesh on that bone. The main body of a horse is typically modeled as a single rigid object. In this model, a horse is a collection of rigid parts connected with hinges. This neglects deformations, or, rather, assumes that the mechanical consequences of non-linkage-like deformations are small enough in their effects so as to be negligible. The key neglected motions are the motion of flesh relative to bone, and that real joints are not precisely hinges nor ideal ball-in-socket joints.

The torques (moments) transmitted across a real horse’s joints can be from various sources:

1. Joint friction, which is presumed negligible for our simple analysis;
2. Complex joint contact wherein there is not a single (*e.g.*, center-of-sphere) effective hinge location, which we do not consider here; we assume frictionless pin joints;
3. Tension in muscles and tendons. These are of central concern for the stability calculations but are (unless otherwise stated) neglected in the equilibrium calculations;
4. Tension in ligaments, which we consider as part of the joint mechanism and do not consider in any explicit way; and
5. Other soft tissue stresses (*e.g.*, skin tension), which we assume are negligible.

This model analysis is a study of 3: the forces and moments due to muscles and tendons (these are neglected in the statics analysis, they are teh topic of interest in the stability analysis).

### Free Body Diagrams (FBDs)

A free body diagram (Fig. 2) is a sketch of the system and of all of the external (from outside the system) forces and moments (torques) on it. When considering equilibrium of a horse we can consider the horse as a whole, or in parts. Here, we will consider free body diagrams of these subsystems:

1. the whole horse including body, head, tail and legs;
2. the horse body, from the legs up;
3. the fore-legs (for a two-dimensional analysis, effectively a single fore-leg); and
4. the rear legs (also treated, in 2D, as a single effective rear leg)..

Forces and moments that are internal to the subsystem do not show on free-body diagrams. For example, in the free-body diagram of the whole horse (Fig. 2), there are no muscle forces shown, because they act internally to that system (or, if you like, they act in action-reaction pairs that cancel each other out). The relevant external forces are, depending on the free-body-diagram,

1. the force of gravity on that part;
2. the force of the ground on the horse feet (for the FBD of the horse and for the FBDs of the legs);
3. the force transmitted across the shoulder or hip joint, from legs to body, or *vice versa*, for free body diagrams of the horse body or of the horse legs;
4. the muscle, tendon and ligament torques at the shoulder or hip, from legs to body, or *vice versa*, for free body diagrams of the horse body or of the horse legs.

In the statics calculations we are finding minimum-effort postures. We seek postures in which the muscle and tendon moments are zero.

### Statics *vs* Dynamics

Outside of discussions of postural stability, a standing horse is generally considered to be standing ‘still’, which means that any motion is negligible. However, all standing involves slight swaying movements due to involuntary functions like breathing. These movements cause motion of the person or animal’s center of mass. Further, using neurologic control to maintain stability involves corrective motions. In other words, a standing horse or human is always moving at least a little bit. For our purposes here, statics (how forces balance when there is no motion) is a good-enough approximation to the more-realistic and more complete dynamics analysis (how forces do not balance, and the consequent motion).

How can you tell whether statics can be an accurate approximation of dynamics? Imagine writing all of the full dynamics equations (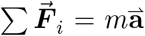, etc.). Then, take those same equations, but set all of the accelerations to zero. Those are the statics equations (Eqs. 5, in Appendix B). If the solutions of those two sets of equations, the dynamics equations and the statics equations, are negligibly different, then one is in a situation where statics can be applied. Generally, this holds true when ‘inertial forces’ 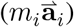 are negligible. That is, when inertial terms are much less than typical forces.

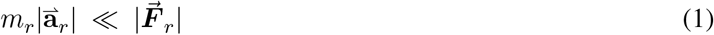

where 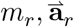 & 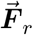 are the mass, acceleration and relevant force, respectively, of the relevant horse part (subscript ‘r’= “relevant”). That is, even though everything is always moving, at least a little, a statics analysis is appropriate if the various forces tend to cancel each other out rather than summing to cause motion. The amount that the foces don’t exactly cancel (meaning that they don’t exactly satisfy the laws of statics, Eqs. 5) gives the little bits of motion. Yet, these little bits of motion, the slight swaying movements of a horse, might be considered negligible for calculating forces. In particular, a horse that is standing and swaying slowly, is probably accurately-enough (at least for study of the choice of basic posture), modeled as always being in static equilibrium without considering the dynamics of swaying. Eqs. 5 will give almost identical answers as would a full ‘inverse-dynamics’ analysis. These calculations are shown below

Unless one is, for instance, studying the leg while kicking, for a horse, typical forces come from the weight *m_horse_g*, where *g* ≈ 10m/s^2^ is the gravitational constant. And inertial forces are associated with the horse accelerations *m_horse_ a_horse_*.

### Estimating the validity of the statics approximation for a standing swaying horse

For a horse swaying back and forth a distance *d*, with frequency *f* (oscillations/s), the acceleration is about *a_horse_* = (*d*/2)4*π*^2^*f*^2^. For example, consider a horse swaying back and forth 1 cm(*d* = .005 m) every 4 seconds (*f* = .25/ s) ([6]).

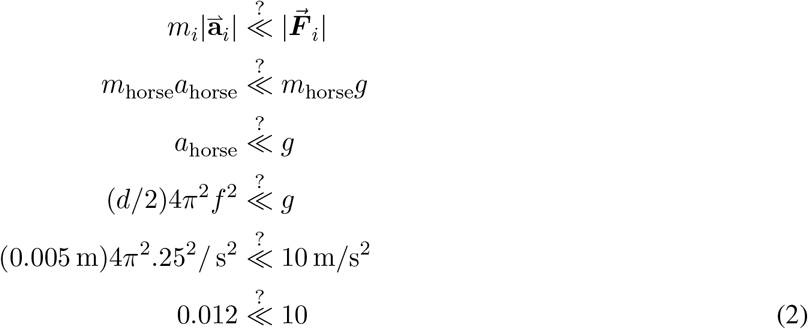

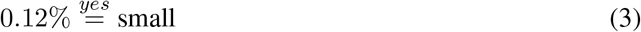

That is, in this case, we make about 0.1% error in force calculations, at least leg compression force calculations, by using statics instead of dynamics. And, for most biomechanics purposes, 0.1% is *much* smaller than other errors we make in muscle modeling, geometry, *etc*. So, by this calculation, statics seems reasonable, at least for calculation of the overall forces involved in holding an equilibrium posture.

Note, however that the calculation above is misleading because the forces accurately calculated with statics are of the legs supporting the horse, whereas the acceleration is of the body’s horizontal (fore-aft) position. If we are interested in the lateral corrective forces, those match the lateral accelerations, and we probably need to use dynamics to understand the motions near the equilibrium positions, and thus we need dynamics to study the stability of these configurations.

Note: there are conceivable situations where study of oscillations near equilibrium could be essentially non-dynamic, meaning non-inertial. For example: maintaining equilibrium in the presence of breathing; or motions due to complex ‘history-dependent’ muscle properties (analogously, some friction-induced oscillations can be understood with just a statics analysis (*e.g.*, [23]). However, if our only concern is whether an equilibrium posture is stable, or not, there is a relevant shortcut that finesses use of dynamics models: using potential energy.

### Potential energy and dynamic stability

If one just wants to know if an equilibrium configuration of a mechanical system (say a posture) is stable, and if the forces can be modeled as either state dependent — meaning that all gravity and muscle forces only depend on the instantaneous positions of the horse parts, with no delays — or are viscous damping terms, a statics analysis is still useful. With such forces, a statics based potential-energy calculation correctly determines whether a configuration is dynamically stable or not (*i.e.*, the system is stable if the potential energy is at a local minimum).

For example, motions near equilibrium of a multi-degree-of-freedom system which are governed by equations of this form:

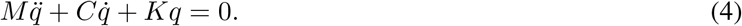

In this equation *q* is a list of numbers describing the deviation from equilibrium and *M*, *C* and *K* are constant (mass, damping and stiffness) matrices that come from linearizing the equations of motion near the equilibrium. Assume that the mass matrix *M* and damping matrix *C* are symmetric. If the stiffness matrix *K* is not symmetric, the system has no potential energy (*i.e.*, the position dependent forces are not conservative). Assuming *M, C* and *K* are all symmetric, the system is stable if the dissipation is always positive (*i.e., C* is positive definite) and if the associated potential energy is always positive (K is positive definite). That is, assuming positive dissipation, the system is dynamically stable if the equilibrium is at the bottom of a potential energy bowl.

In our case, with only one degree of freedom, *q* is just a single scalar number (say, the deviation of the hip joint angle from equilibrium), and *M, C* and *K* are also numbers. Assuming *C* is positive (damping is dissipative), stability is determined by whether *K* > 0 (stable) or *K* < 0 (unstable). That is, dynamic stability can be determined by calculations of potential energy in the neighborhood of the equilibrium (a statics calculation of sorts) while never doing a dynamics calculation. In particular, for our horse model, we determine dynamic stability by looking at the net potential energy due to gravity (which contributes to making K negative) and due to the corrective springs (which contribute to making potential energy positive).

There is a separate question (discussed in Appendix E, below) whether this potential energy approach is a valid approximation if there are time delays.

### 2D vs 3D

Here, we are doing a two-dimensional (2D) analysis. We view the horse from the side, considering only an analysis in the *sagittal = side-view = lateral-view* plane. In effect, then, the horse has two legs: the front leg(s), representing the pair of fore-limbs(s) and the back leg(s), representing the hind-limbs. We use a 2D, instead of 3D, analysis because:

1. The 2D results are simpler to understand than 3D results, in particular, the fewer the number of forces to consider, the more-digestible are the results;
2. A real horse may be decently modeled as 2D some times and for some purposes:
  - The side-to-side (left-right, medio-lateral) forces can be relatively smaller than fore-aft (cranio-caudal) forces;
  - The left-right (coronal, frontal plane, front view) leg splay is relatively smaller; and
  - The horse and its posture are often nearly left-right symmetric (so side-to-side forces cancel); and
  - The 2D results are exact features of the 3D mechanics; that is, if all 3D forces are projected into the sagittal (side view) plane, and the forces on all horse parts are considered as the set on the left & right pair added (*e.g.*, the rear legs ground force is the sum of all forces on both rear feet, added up), then the 2D statics results are exact results for the projections of the 3D forces and moments.

## Appendix B: The horse model

In this appendix we consider some details of the horse model. The horse (Figure 2 on page 3 and Figure 4 on page 21) is made of three rigid objects: 1) the fore-legs with length *ℓ_f_*; 2) the rear legs with length *ℓ_r_*; and 3) a back with length connecting the front hoof F with the shoulder S and the rear hoof R with the hip H, respectively. The hoof ‘points’ F and R are at the centers of pressure of the hoof ground reaction forces (*i.e.*, the points where the equivalent single ground-reaction force-couple systems have no net torque). We do not attempt to find this point accurately, and take it to be at the rough ‘middle’ of the footprint. We neglect the weights of the legs. The horse mass is all attributed to the back and head. The center of mass G is a distance *d_G_* back from S along SH, and *h_G_* orthogonally above the line SH. The distance between the legs, from F to R, is *ℓ* = *d_f_* + *d_r_*, where *d_f_* and *d_r_* are the distance from the ground-projection of the Center of Mass G to the front and rear hooves, respectively.

**Figure 4:**
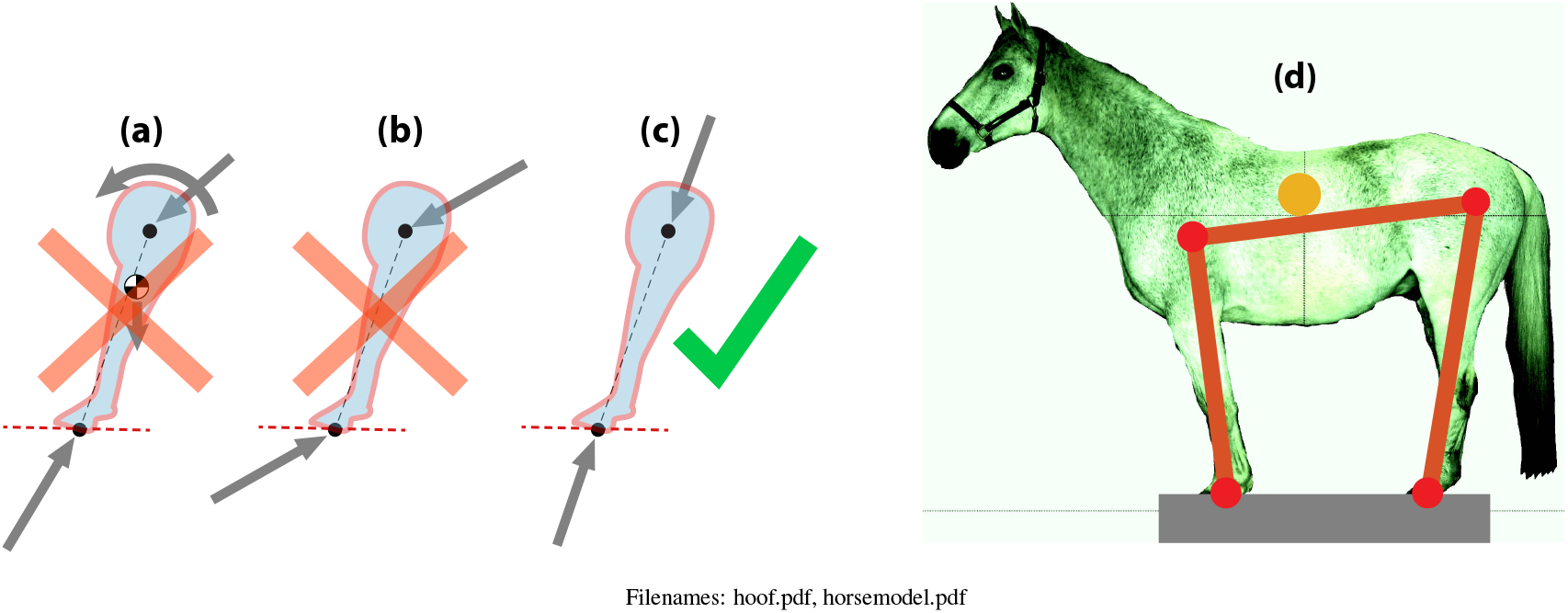
**Leg as a ‘two-force object’** and **Horse model.** If we neglect the weight of the legs, the legs are two-force objects. **a) General case.** This free body diagram includes the leg weight and also torques from muscles at the hip. **b) Two forces,** mechanics reasoning not yet applied. This free body diagram leaves off the leg weight and also torques from muscles at the shoulder or hip, but mechanics reasoning has not yet been applied. As drawn, it is not in equilibrium. **c) The leg as a two-force body.** Leaving off the hip torque and the leg weight necessitates that the two forces be equal and opposite and along the line connecting the points of application. This is the assumption we use in this paper. **d) The horse is made of three links:** forelegs, back (hind) legs; each pair is one link in the model. These 3 parts are connected with two hinges at the shoulder and hip.

### A standing horse is a three-force object

First, consider the horse as a single stationary object with no concern for the presence, or absence, of muscular effort. We use 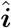 to indicate the direction to the right, towards the horses rear, and 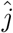 to indicate vertical upwards. There are three forces acting on a whole horse considered as a single object:

1. The force of gravity,

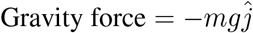

acting at the center of mass G (or CoM) of the horse;
2. The total force of the ground on the front hooves (Fig. 4 on page 21), 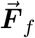 which can be decomposed in the vertical *N_f_* and fictional (horizontal) part *H_f_*, with

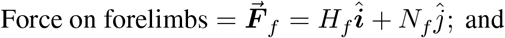
3. The total force of the ground on the rear hooves, 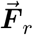 which can be decomposed in the vertical *N_r_* and fictional (horizontal) part *H_r_*

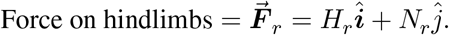

Neglecting the small motions during standing, and neglecting small forces (*e.g.*, wind), we can apply the laws of statics, using only the two ground forces 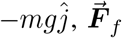 and gravity 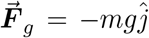, three forces in total. Hence the horse is a so-called ‘three-force object’.

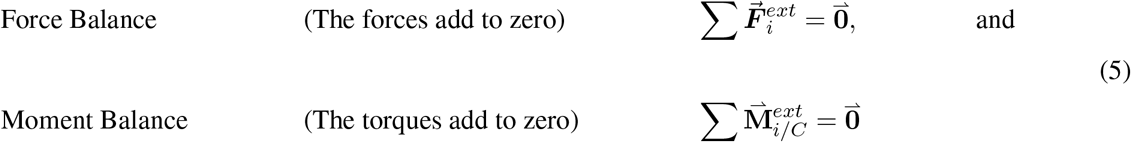

The moment balance (or torque balance) equation holds relative to any point C. As per any introductory statics book (*e.g.*, [24]), we can use these two principles to draw various conclusions about the forces.

### Features of 3-force object equilibrium

1. **‘3-force object’.** Consider a point C at the intersection (if such exists) of the lines of action of the forces at the front hoof and at the rear hoof. The only force with moment about C is gravity. For there to be no net moment the gravity moment must also be zero. So, moment balance about C implies that the two ground reaction forces, at the forelimbs and hind-limbs, are either (Fig.2 on page 3)
  i. **vertical**, or
  ii. have **lines of action that intersect at a point directly above G**, or
  iii. have **lines of action that intersect directly below G**;
2. **The ground carries the weight.** The vertical component of force equilibrium implies that the vertical ground reaction forces add up to the weight: *N_f_* + *N_r_* = *mg*;
3. **Lever rule.** Moment balance about the point on the ground under the center of mass implies that *N_f_ d_f_* = *N_r_ d_r_*
4. **Vertical forces are determinate:** Moment balance about the rear feet and about the front feet, together imply that the vertical ground reaction components are: *N_f_* = (*d_r_/ℓ*)*mg* and *N_r_* = (*d_f_/ℓ*)*mg*;
5. **Horizontal forces cancel but are indeterminate.** Force balance in the horizontal 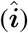 direction implies that the horizontal ground reaction forces are equal and opposite, *H_f_* = −*H_r_*. These forces are otherwise indeterminate (*i.e.*, the magnitude of these canceling forces cannot be found from the laws of statics);

Any 2D statics model of a horse, and any measurement of a real horse that is effectively stationary, must have forces obeying all of the above relations. These apply no matter whether the leg weight is included, or not, and whether or not there are muscle contractions making the legs not ‘two-force-objects’. These, above, are the strongest, most model-independent, results from statics as applied to whole horses in the sagittal plane.

### Approximating the legs as two-force objects

Treating the horse as a 2D linkage with left-right symmetry in structure and in coordination, we can consider the two forelimbs as a single object which we will call the fore-leg, and similarly for the hindlimbs. Before simplifying, we show a free body diagram of a horse leg in Fig. 4a on page 21 which shows these forces:

- the ground reaction force 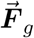 acting on the hoof at the ground. This ground reaction force 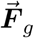 includes both a normal (vertical) component and a sideways frictional component;
- the gravity force on the leg *mg* pointing down;
- the joint force, from the body to the leg, at the hip 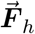; and
- *M_h_*, the moment of the hip muscles, tendons and ligaments at the hip. These act on the leg from the horse body.

Because horse mass is much greater than the leg mass, we neglect the leg mass. Because we might assume that the horse chooses standing postures which minimize muscle tensions and thus minimizes joint torques, we neglect torque at the joints (for finding equilibrium postures here, the muscle forces are central in the stability calculations). Thus, the free body diagram of a leg (pair) has on it only two forces: one from the ground and one from the shoulder (or hip) as shown in Fig. 4b. These forces can only be in equilibrium if the forces are equal and opposite and have a common line of action (’two-force members’ or ‘two-force bodies’ in *e.g.*, [24]), as shown in Fig. 4c.

> *The forces on a horses leg, at the hip and ground, are well approximated as being equal and opposite and along the leg (along the line connecting the hip and hoof).*

So, the ground reaction force at the front hoof points towards the shoulder and the ground reaction force at the rear hoof points towards the hip. This modifies Fig. 2
 to include information about the leg geometry (Fig. 4c on page 21).

### Equilibrium of standing, taking account of the legs as two-force objects

Taking account that the legs are two-force objects changes the results of Fig. 2 on page 3 to those shown in Fig. 4 on 21. That is, in addition to the hoof forces intersecting at a point above or below the center of mass, they are along the lines from the hooves to the shoulder and hip. For a pendulum we could recognize equilibrium by the center of mass being above or below the hinge. Whereas, for a horse:

> *The equilibrium posture of a horse has the lines along the legs intersecting directly above or below the horse center of mass.*

The point of intersection of the two leg lines is, for a horse, somewhat analogous to the location of the hinge for a pendulum; in both cases, the center of mass has to be directly above or below that point in order to achieve equilibrium (see Appendix D for further discussion of the pendulum analogy).

## Appendix C: Calculation of stability of a standing horse

These calculations are the same in spirit as the similar calculations for an inverted pendulum shown in Appendix D. Appendix D, below, on the inverted pendulum analogy, should be read first if this calculation is hard to follow. The standing horse can be considered as a 3-object mechanism (Fig. 4 on page 21).

1. The fore-limbs are hinged to the ground at the front hoof (hooves) and to the body at the shoulder;
2. The body is hinged to the forelimbs at the shoulder and at the hip; and
3. The rear legs are hinged to the horse at the hips and the ground at the rear hoof (hooves).

Note: in mechanical design, this 3-link model is called a ‘four-bar-linkage’ (the ground is the so-called fourth link). Given the position of the hooves and the lengths of the body parts, this linkage has one degree of freedom. That is, one number, say the angle *θ_f_* of the fore-legs, determines the all of the other positions and angles.

### Calculation of stability using potential energy

At each configuration of the linkage there is a gravitational potential energy *E_p_*, which is the weight times the height of the center of gravity,

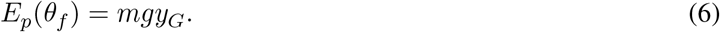

Note, to use this equation one must find the height *y_G_* of the center of mass in terms of *θ_f_* (for the horse mechanism, this is a non-trivial geometry/trigonometry problem with no simple formula). Any equilibrium posture (linkage configuration) is one in which all of the laws of statics hold for the whole horse and any of its parts. Assuming legs with negligible mass and negligible muscle and ligament torques, the possibilities are shown in Figure 4 on page 21. A theorem from statics is that such a configuration is also one where the potential energy is ‘stationary’, meaning,

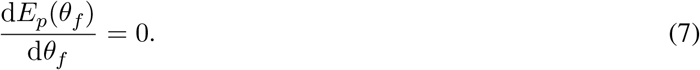

This means that the equilibrium (zero muscle force) posture of a horse is at a (local) potential energy maximum, or minimum, or a stationary point. In all of our calculations for semi-realistic horse proportions, we always find the equilibrium postures to be at a local maximum of potential energy; the energy function is an upside-down bowl. That is, we always find that when we find a *θ_f_* that satisfies Eqn. 7 that it also satisfies

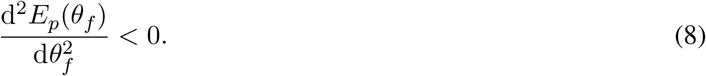

Such an equilibrium, at a potential energy maximum, is unstable. If disturbed ever-so-slightly from this configuration the horse would, if there were no ligaments, muscle forces or other joint torques, fall down. That is, in our three-link model, a standing horse in equilibrium is, with no muscle effort, unstable. Nonetheless, horses do stand for extended time without falling. So, horses must apply corrective torques. That is, when the horse is slightly out of position (slightly away from the zero-torque equilibrium posture), there must be corrective torques that bring the horse back to, we presume, near the zero-torque equilibrium posture. There are a variety of known sources of these torques, namely

1. Muscular forces actively (dynamically) controlled using sensory and motor loops;
2. Muscular tonic contractions, where the intrinsic stiffness of active muscles resists joint-angle changes;
3. Passive ligaments that stabilize the joints; or
4. A bone and ligament morphology that effectively locks the joints (*e.g.*, the equine check and stay apparatus).

All of the mechanisms above likely have the feature that deviations from equilibrium posture lead to corrective torques, thus the behavior of any or all of them is somewhat like that of torsional springs, *k_f_* and *k_r_*, at the joints. So, we now consider joint torques that only apply if the joint angles, *θ_f_* and *θ_r_*, deviate from equilibrium, these are:

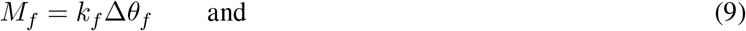

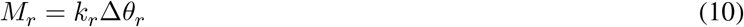

where Δ*θ_f_* and Δ*θ_r_* are the deviations of the joint angles relative to equilibrium. The addition of these springs does not change the equilibrium posture, but the springs do change the stability. Namely, the potential energy is now:

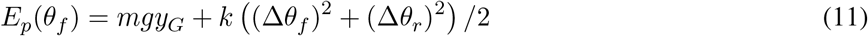

where, for simplicity of modeling, we use the same spring constant at both joints, *i.e., k* = *k_f_* = *k_r_*. For a given horse model with given hoof placement we can solve for the equilibrium posture (configuration). And, using Eqn. 11 we can also find the minimum value *k_min_* of *k* in order to make the posture stable. That is, we apply Eqn. 11 to the critical condition (inequality in Eqn. 8 turned to equality),

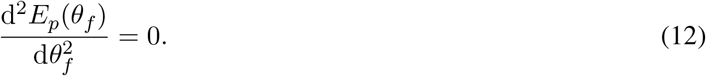

and solve for *k*, calling that *k_min_.*

For each hoof placement, canted-in, vertical, or splayed-out, we can find the associated equilibrium posture one of two ways: either using the two-force-object and three-force-object reasoning above, or by finding stationary points for the gravitational potential energy. As described so far, the model allows some non-physical postures, for example, hanging upside down from the ground surface. We exclude this. Depending on leg and body lengths, there can also be very-crooked equilibrium postures, with the hip much above or much below the shoulder. And postures above and below ground with the legs crossed and the body possibly upside down. There is just one above-ground posture that has the legs not crossed, the body near level and that satisfies the equilibrium equations. Our numerical calculations find that this posture is always unstable. For this one equilibrium above-ground posture, we seek the minimum needed stiffness *k_min_* for it to gain stability. Because of the complex geometry of the linkage, and lack of symmetry, both the find-equilibrium and evaluate-stability calculations are done numerically.

### Numerical calculations

The horse length and mass parameters are given in Appendix F. We chose to parameterize the postural choice with leg splay as measure by the hoof-to-hoof ground spacing, the length *ℓ_g_*.

#### Find equilibrium geometry using numerical root finding

Given *ℓ_g_* we then consider a candidate value for the slope *θ_f_* of the fore-leg. Given *θ_f_*, the law of sines and the law of cosines finds the full 4-bar linkage geometry, including all link angles and all joint positions. We then did numerical root finding to find that value of *θ_f_* which put the intersection of the two leg lines directly above, or directly below, the horse center of mass. That configuration is, as per the three-force-object reasoning, the equilibrium standing configuration for the given leg splay.

#### Numerical stability calculation

Given the equilibrium configuration for given *ℓ_g_*, we then found the equilibrium configuration’s stability as follows. We varied *θ_f_* in the neighborhood of the equilibrium position. For each *θ_f_* we found a consistent linkage geometry and associated center of mass height and hence (multiplying by mg) potential energy. We then did a polynomial fit of that function and recorded the quadratic term. We similarly found the best fit for the sums of squares of the shoulder and hip joints *vs θf* in the neighborhood of equilibrium. Given those two fits we could find that torsional spring constant k so that if one such spring was put at the shoulder, and the other at the hip, that the net quadratic term in the potential energy would be zero. This is the so so called “minimum required joint stiffness” *k*_min_, displayed graphically in Fig. 3 on page 8.

### Intuitive explanation of mechanics

The basic result, that a splayed-out posture is more stable (by our measure), can be predicted qualitatively without need for detailed calculations. For the torque at a joint to have effect on the whole mechanism motion, the whole-mechanism motion must involve changes in that joint angle. Imagine a horse that is so canted-in that the front and rear hooves nearly touch. Then, if that horse rocks forward (or backward), still keeping its feet on the ground, there is significant falling even though there almost no change in the shoulder or hip angles (the horse falls almost like a triangle rocking on its lower vertex). Thus, for that extremely canted-in posture, springs at the shoulder or hip joints would have almost no effect (have almost no righting torque, cause almost no change in potential energy, do almost no work). When the legs are more spread, the joint angle changes are bigger for a given rocking. Bigger joint-angle changes have a dual effect: bigger angle changes with a given amount of falling cause bigger rotation-induced spring torques; and, due to the reciprocal nature of mechanical advantage, also, have a bigger mechanical advantage (*i.e.*, bigger effect on the mechanism for given spring torque). Meanwhile, the curvature of the gravitational potential energy near equilibrium, as a function of leg splay, is slight (the circular arc of CoM motion when the legs are parallel is similar to the near-circular arc when the legs fully canted-in). So, due to the enhanced utility of the springs in splayed-out posture, the splayed out posture is ‘more’ stable (by our measure, which is *not* the only plausible measure, see appendices D and E below).

## Appendix D: Pendulum analogy

The concepts of equilibrium, and stability of equilibrium, are perhaps best understood using a simpler system. Consider a pendulum: a stick with length *ℓ* connected to the fixed ground by a hinge at H (Fig. 5a on page 28). The stick, while not looking like a horse, exemplifies the concepts of equilibrium and stability of equilibrium. The average position of the stick mass is at the middle G of the stick. This is the center of mass (CoM). The hinge H is a distance *d* from the lower end of the stick at A. The hinge H is assumed to be well made and well lubricated, so does not resist rotation of the stick. However, we could apply torques at the hinge with muscles, or some proxy for muscles (*e.g.*, a motor or a torsional spring). In this analogy, ‘posture’ is the chosen position of the hinge *d*, and the chosen angle *θ*. Now consider equilibrium, and stability of equilibrium, of this stick considered as functions of the parameters.

**Figure 5:**
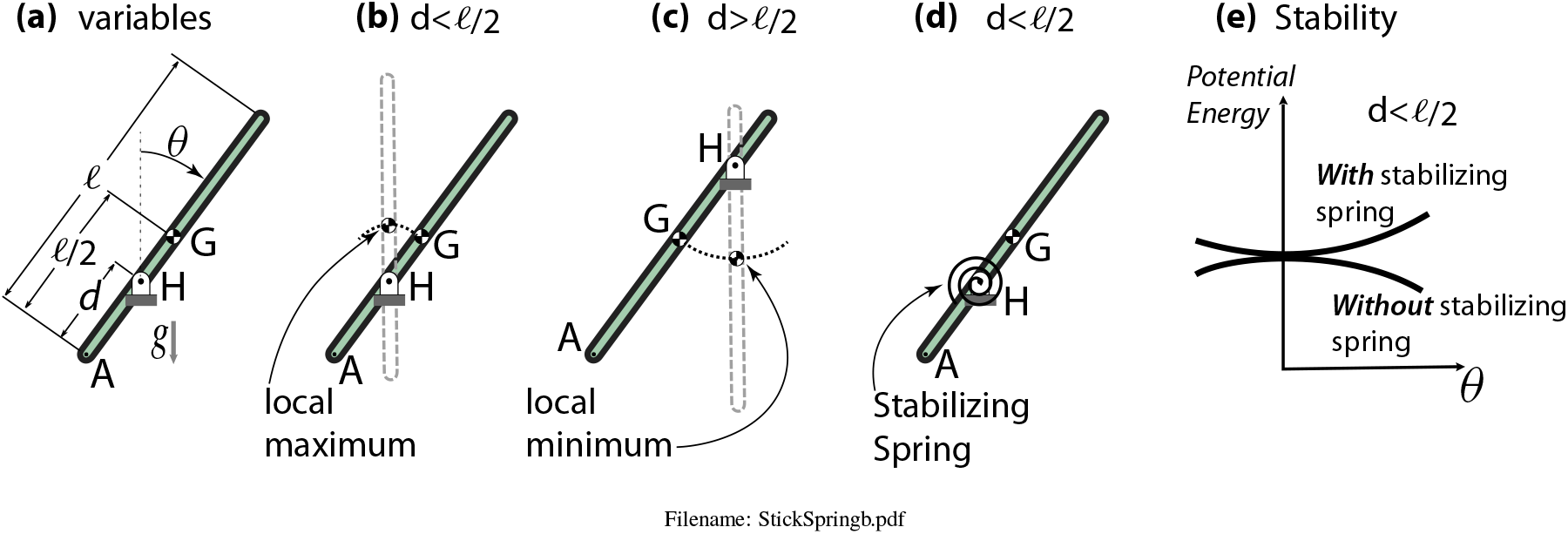
Pendulum: Stability, and stability of equilibrium. **(a)** End at A, hinge at H, Center of Mass at G, tipped an angle *θ* from vertical; **(b)** Center of mass above the hinge is unstable, equilibrium is at a potential energy maximum; **(c)** Center of mass below the hinge is stable, equilibrium is at a potential energy minimum **(d)** A spring is added to the unstable configuration; **(e)** The unstable configuration is made stable by the addition of a spring, changing a potential energy maximum (bowl down, unstable) into a potential energy minimum (bowl up, stable).

### Equilibrium ‘posture’ for a pendulum

All of the configurations drawn in Fig. 5 on 28 have *θ* ≠ 0 and *θ* ≠ *π*, they are all non-equilibrium. Equilibrium configurations are shown dotted in Figs. 5b and 5c. If the stick is put in a *non*-upright (non-equilibrium) configuration and the hinge is not at the center of mass (*d* = *ℓ*/2) then, if it is released from rest it will tend towards an upright position: If the center of mass G is above the hinge (*ℓ*/2 > *d*, Fig. 5b), it will fall away from upright; and if the the center of mass is below the hinge (*ℓ*/2 < *d*, Fig. 5c) it will swing up towards vertical. Thus, all tipped positions are, unless the hinge is *at* the center of mass (*d* = *ℓ*/2), not equilibrium positions; the moments do not balance and the stick tends towards upright, by falling or swinging. The set of equilibrium positions are places where (a perfect) stick can stay indefinitely without falling. The equilibrium positions are those for which either the hinge is at the center of mass (*d* = *ℓ*/2), or for which the stick is upright (*θ* = 0 or *θ* = *π*).

Assuming no disturbances, it takes no torque to hold an equilibrium position. This is true if H is above G (*d* > *ℓ*/2) or if H is below G (*d* < *ℓ*/2). Alternatively, the stick stays in place if *d* = l/2. In short, the set of equilibrium positions are then *θ* = 0, for all values of *d* and for *d* = *ℓ*/2 for all values of *θ*. (The special case when G is at H and there are multiple equilibrium orientations, is not relevant to the always-unstable horse discussion.)

### Stability of equilibrium

The vertical equilibrium postures are, intuitively, quite different depending on whether the center of mass G is above (*d* < *ℓ*/2, Fig. 5b) or below (*d* > *ℓ*/2, Fig. 5c) the hinge. Actually, if the center of mass is above the hinge, many people have trouble accepting that that is indeed an equilibrium position at all. If the center of mass is above the hinge, the stick would obviously fall. But ideally, if it was a perfect stick, and placed perfectly vertically, it would not fall, at least not for a very long time. So the position where the center of mass is directly above the hinge is still, at least in the language of mechanics, ‘in equilibrium’; to keep that stick upright you don’t need to apply a torque to the left, nor to the right. On the other hand, practically speaking, you have to hold it to keep it from falling. That is, the upright configuration is an *equilibrium* posture, but it is an *unstable equilibrium.*

- **Equilibrium** postures can be held with no extra torque at the hinge.
- **Non-equilibrium** postures require constant joint torques to be held.
- **Stable equilibrium** postures are ones where, if disturbed from equilibrium a tiny amount, the system will tend to return towards that equilibrium with no muscular torques.
- **Unstable equilibrium** postures are ones where, if disturbed from equilibrium a tiny amount, the system will tend to exponentially deviate from the equilibrium.

#### Potential energy

The stick is in equilibrium if it is vertical. The vertical equilibrium is only stable if the center of mass hangs below the hinge (*d* < *ℓ*/2, Fig. 5c). The vertical equilibrium is unstable if the center of mass is above the hinge (*d* > *ℓ*/2, Fig. 5b). More technically, the equilibrium is stable if the system potential energy is at a local minimum, and unstable if it is at a local maximum (see Fig. 5e). For this system, the potential energy is proportional to the height of the center of mass. Relative to nearby configurations (nearby values of *θ*) the vertical stick is at a locally minimum-energy height (*i.e.*, stable) if the center of mass G is below the hinge H. And it is at a locally maximum height (*i.e.*, unstable) if G is above H.

### Quantifying pendulum instability

In relation to the more complex horse problem, the relevant pendulum cases are when the hinge is below the center of mass (*d* < *ℓ*/2); having *d* < *ℓ*/2 is like the horse being above the ground; having *d* > *ℓ*/2 would be analogous to a horse hanging by its hooves upside down from the ceiling 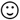 (to be complete, there are also non-physical cockeyed equilibrium postures that our horse model can exhibit, but which a real horse could not exhibit). We are only interested in intrinsically unstable equilibrium postures. These are analogous to an ‘inverted’ pendulum.

We can make an unstable posture into a stable posture by always being at the ready with corrective torques. For example, if we add a stiff-enough spring to the hinge, one that tends to center the stick at *θ* = 0, we can make an unstable stick into a stable stick (Fig. 5d). One way of quantifying the amount of intrinsic (pre-correction) instability is by the size of the spring needed to make the unstable configuration stable. For the pendulum, the total potential energy is

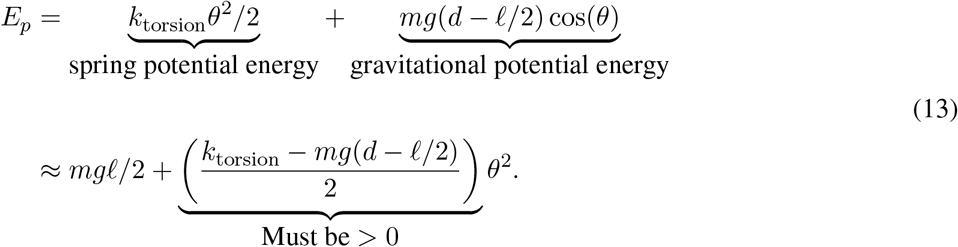

We used the small-angle approximation that cos *θ* ≈ 1 – *θ*^2^/2. To make *E_p_* an upwards shaped bowl function of *θ* (that is, to make the equilibrium a potential energy minumum) we need the coefficient of *θ*^2^ to be positive (see Fig. 5e). So, stability only occurs if the torsional spring has a big-enough constant

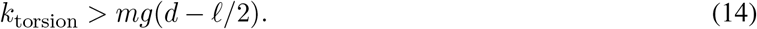

The otherwise-unstable vertical pendulum is stabilized by the addition of a *k*_torsion_ satisfying the inequality above.

Adding the (big-enough) spring potential energy to the gravitational potential energy changes the upright configuration from a potential energy maximum (dynamically unstable) into a potential energy minimum (dynamically stable). With a big enough spring, as *θ* is deviated from 0, the increase in spring energy will win over the decrease in gravitational potential energy. The more that the center of mass is above the hinge (the bigger the distance *d* – *ℓ*/2), the bigger the needed stabilizing spring needed to make this otherwise unstable equilibrium posture stable.

A spring is equivalent to a linear (proportional) controller. So, our measuring uncontrolled instability by the minimum added spring to gain stability is like measuring the amount of uncontrolled instability by the minimum feedback gain needed to obtain stabilization. Thus, we might think of this spring as a proxy for the amount of neuro-muscular control needed to hold a pendulum upright. This neuro-muscular control could be through a feedback loop or through tonic muscular action, such as with co-contraction. In a feedback loop, the deviation would be sensed, and then corrective muscle contractions commanded. In, say, co-contraction, pairs of opposing muscles would be continuously held on, and the pair of muscles would act like a corrective spring. In either mode of control, the bigger the needed spring in the model, the bigger the needed neuro-muscular control in the biological system.

With the horse model, the situation is analogous. The calculations for the horse model are more complicated, and in the horse model, the amount of spring needed depends on the posture (the spacing between the feet).

#### An alternative quantification of instability

**eigenvalue.** If disturbed slightly from an equilibrium configuration, an unstable system diverges exponentially from the equilibrium. An alternative quantification of instability to our *k*_min_ described above, would be the characteristic time of this exponential growth; the shorter that time, the faster the growth, the more unstable is the system. The inverse of this characteristic time is the ‘eigenvalue’ λ (meaning, characteristic value). By the eigenvalue measure of instability, the bigger the eigenvalue the more unstable the system. More technically, near an unstable equilibrium all quantities grow in time in proportion to *e*^λt^, so λ measures teh rate of growth.

Consider a point-mass inverted pendulum with massless stick length *ℓ*, point mass *m* and gravity *g*. Now consider a variety of other pendula with different values for length, mass and gravity, and compare their instability by the two measures above (*k*_min_ and eigenvalue λ.

1. Double *m*. This doubles the needed stabilizing spring *k*_min_ but has no effect on the eigenvalue λ. Thus, this would be a more unstable pendulum by the *k*_min_ measure and equally unstable by the λ measure.
2. Double *ℓ*. This slows the characteristic time of falling but increases the spring needed for stabilization. That is, the two measures of instability have opposite trends for pendulum length changes.

#### Generalized measure of instability

Imagine balancing an upright broom on the palm of your hand by moving your hand around. Now imagine trying to similarly balance a pencil. Clearly, in that context one would say that a pencil is more unstable because it is harder for a person to balance. On the other hand, imagine an upright 5m (quite tall) ladder on the ground, but leaning against nothing. You are to hold it there. Then imagine a person climbing to the top, and you are still trying to balance the ladder. Clearly the ladder is more unstable when there is a person on top. For the pencil vs broomstick, the eigenvalue measure seems a more appropriate measure of instability. For the ladder vs ladder with person up top, the minimum-stabilizing-spring is a better measure.

What is a generalization that uses the right measure in the right context? One idea is to measure instability by the difficulty of control using the hardware and software that are available for actual control system that is in use. The pencil is more unstable than the broomstick because it demands a neural control loop that is faster than a person has available. The ladder with a person up top is more unstable because, for the size of deviations from vertical a person can detect, the heavy-topped ladder needs more strength than the person has available.

If one knows which issue is the main challenge for the control system of interest, one can pick an appropriate measure of system instability. If speed of response is the critical issue, the eigenvalue measure is more relevant. If size of the required force for a given deviation is critical, then the minimal-stabilizing-spring, our choice for this paper, is most relevant.

## Appendix E. Neglect, or not, of neural delays; passive tonic vs active feedback

### Corrective torques

As noted, a horse’s standing posture is unstable without corrective torques. There are various possible sources for these torques listed in Appendix A. We focus on the corrective torques from muscles (more precisely, muscle-tendon complexes). These could be of three kinds:

1. **Tonic stiffness.** Muscle torques based on muscle’s intrinsic stiffness. This is the slope of the muscles force-length curve in a quick (too fast for neural feedback) imposed extension (neglecting, or subtracting out, viscous/hystoretic effects). Intrinsic stiffness does not involve feedback via neural control. As noted in *e.g.*, [27] muscle intrinsic stiffness is more-or-less proportional to the present muscle force. So, intrinsic stiffness is only significant if a muscle is activated. There are two ways this can happen in a static posture:

a. Co-contraction, with flexors and extensors in opposition; and
b. Contraction opposing an external force. For example, this non-co-contraction tonic state is generally thought to be an approximation of the state of a typical standing person (*e.g.*, [19]); the person is leaned forwards of equilibrium to put their center of mass slightly forwards of the ankle, with gravity causing a disruptive torque (about the ankle hinge) and the achilles tendon (with soleus and/or gastrochnemius muscles) providing a counter-acting righting torque.

One might term the stabilization from intrinsic stiffness as ‘tonic’ (as due to a constant activation), ‘passive’ (as not involving active neural control), or even as a primitive form of ‘feed *forward*’ (in that it is a chosen activation pattern, albeit simple, that achieves the desired coordination without feed*back*);
2. **Corrective muscle torques** due to sensed deviations from equilibrium. These would be based on proprioception (mechanoreceptors distributed in muscles, joints, ligaments and skin), visual (oculo-vestibular responses), or inner ear (vestibular end organs– otoliths and semicircular canals), processed in the spine or brain, and then leading to muscle commands. This corrective strategy might be termed as ‘feedback’ (activation based on sensing is fed back to the muscles) or as ‘active’ (in that the neural output is actively changing);
3. **A combination** of the two mechanisms, with some of the corrective torque do to tonic (intrinsic) stiffness and some due to active feedback control.

### Feedback delays

For active neural feedback in an animal (or human) there are delays due to sensor delay, neural transit times to the spine or brain, neural processing time, neural transit time back to the muscle, and muscle delays (due to electrical, chemical and mechanical activation effects). These have a variety of delay times, depending on the pathways, that taken together are thought (for humans) to be in the range of 0.1s - 0.3s in [12].

A given postural control model may concern passive (tonic) stiffness or active (neural feedback) control or both. And the model may take account of neural delays, or may not. Our model here neglects neural delays and assumes proportional feedback. So, in our model, tonic stiffness and active proportional feedback are equivalent and inseparable, they both yield a (modeled as) instantaneous corrective torque proportional to distance from the target state.

### Grasp

The stability of grasp has been studied with a similar philosophy as used in this paper. In both [21] and [27] only tonic (intrinsic, passive, instantaneous) corrective torques were considered. As pointed out in [27], if only the second of these (in the list above) is considered (muscle fighting a load) the stiffness is proportional to the load. Applied to grasp, this muscle model (stiffness proportional to force) says that, without co-contraction, a grasp configuration is either stable, or not, depending on the geometry of the grasp but independent of the load; higher grasp forces proportionally increase both the geometric negative-stiffness and the muscle tonic stiffness. So, assuming muscle stiffness exactly proportional to load, one stiffness is bigger than the other for all grasp forces. If the intrinsic stiffness is greater, the grasp is stable. If the negative pre-stress (geometric) stiffness is greater, the grasp is unstable. (More exactly, for a multi-degree-of-freedom linkage model with this type of muscles, there is a net stiffness which is either positive definite or not, depending on geometry and not on the tightness of the grasp).

For the grasp stability studies mentioned above, feedback loops were not considered. This is reasonable — the characteristic time of grasp instabilities are so fast compared to neural delays, because of the low inertia of fingers, that there is not possibility of stabilization from (delayed) feedback loops.

### Tonic stiffness and standing stability

As just noted, for human grasp, mechanism stability is neither improved nor decremented by the tightness of the grasp. However, applying this muscle-stiffness-is-proportional-to-force muscle description to a standing person or horse shows that stability can be enhanced by standing further away from equilibrium. That is, the effectively-negative gravitational stiffness is nearly constant with distance from equilibrium, but the muscle stiffness increases with distance from equilibrium because the needed muscle force, and thus muscle stiffness, goes up with distance from equilibrium.

### Neglect of intrinsic stiffness in human standing models

In the study of the effect of leg spread on the stability of human standing, [3] ignore the intrinsic stiffness of muscles, only considering active feedback mechanisms that have neural delays. Similarly, [12] focus on neural feedback and not on ‘passive’ (meaning intrinsic without active neural control) stiffness.

Finally, [19] studying human standing claims that there is some stiffening from the intrinsic stiffness of active calf muscles and, on top of that there is a dynamic feedback loop.

It is sensible that an animal or person would choose to stabilize using a mixture of tonic (constantly contracted muscle) and feedback control, for two reasons (Art Kuo — personal communication):

1. There is a cost to varying muscle force. This cost is on top of the cost of maintaining a muscle force. Given that the sensors have limited resolution, feedback control necessarily involves force variation. If some of the stability is handled by tonic contraction (using intrinsic muscle stiffness), then there is less instability for the feedback system to contend with, and, for given sensor thresholds, smaller needed corrective torques. Depending on details of muscle costs for tonic forces vs the costs for the necessarily time-varying contractions used in feedback, there could be a savings of chemical energy used by using a mixture of tonic (intrinsic stiffness) and feedback control.
2. Muscles and tendons are non-linear, they are softer when the force is smaller. Starting with no force, to generate a small force slack needs to be taken up. Whereas, when a muscle is already contracted it takes less change in excitation, and there is less related muscle and tendon stretch, to effect a similar force change. That is, modulating an existent force is easier than modulating a force near to zero force.

### Combining active feedback and passive tonic controls

As noted, our model treats active feedback control and passive tonic control as equivalent. That is, we assume that neural delays are not significant. Consider two animal systems A and B. Assume A can be stabilized with a smaller spring *k*_minA_ than is needed for system B. No matter what the actual mechanism the animal uses, whether it is intrinsic tonic stiffness or feedback loops, or both, we assume that it is easier for the neural system to stabilize system A, with *k*_minA_ < *k*_minB_, than B.

The error that we could be making is that neural delays could be a part of the challenge of postural control. If the observed sway dynamics has characteristic times comparable to delay times, or shorter, then delays probably cannot be neglected. If the characteristic sway dynamics have characteristic times much longer than the neural delay, than the neural delays are probably negligible and feedback loops are essentially dynamically equivalent to springs.

### Quick review of, and comparison with, Bingham *et al* [3]

For analysis of standing humans in the frontal plane, [3] use the same mechanical model as we are using for the horse, but with different parameters (that which is the horse body in our model is the human pelvis in their model). Their two key model findings are:

1. They notice that their model is *less* stable with spread legs than with vertical legs; and
2. They find that there is a range of feedback gains which stabilize the system. This range is smaller for spread legs than for parallel legs.

These results are, at least superficially, in contrast with our model results that

1. Our horse is *more* stable with legs splayed-out than with legs parallel; and
2. The minimum stiffness needed for stabilization is less for a splayed-out horse than for a parallel-legs horse.

Besides the differences in physical parameters (lengths, masses), our model differs from [3] in two key ways:

A. They use the eigenvalue λ as a measure of the degree of instability whereas we use the minimum corrective spring *k*_min_; and
B. They assume all control is with neural feedback that has a delay, whereas, whether tonic or feedback-based, we neglect the delay.

### The differences between this paper and Bingham *et al*

First, they have the result that splayed-out is more *un*stable, in contrast to our finding that splayed-out is more stable. Their result might be explained intuitively by the splayed-out posture being associated with a smaller inertia (less kinetic energy per unit leg angular-rate). Thus, although the gravitational negative stiffness is smaller with splayed out legs, the reduced inertia makes for a reduced characteristic time of falling (bigger eigenvalue) and thus less stability by their measure. On the other hand, when the legs are splayed out, a given motion of the center of mass is associated with large changes in the body-to-leg angles, so that our model’s joint springs have a larger stabilization effect. Thus, by our measure, splayed out is more stable.

Second, is their result that splayed out is harder to control, in contrast to our result that splayed-out is easier to control. As noted just above, when splayed-out, the joint angles change a lot with center of mass motion, so any joint torques have larger control authority. Thus our result, that the minimum stabilizing springs are smaller. And thus, in our interpretation, less neurological control is needed (meaning smaller co-contraction or smaller leaning and mono-tonic contraction or smaller feedback gains). Consistent with us, they find that the minimum gains to stabilize the splayed out posture are smaller. But, their model includes neural delay. If one uses a gain much larger than the minimum-needed gain, then, in their dynamics model, the characteristic sway becomes faster with shorter characteristic times, eventually competing with neural delay times, and the control system becomes unstable (there is an oscillatory instability at a critical gain). On the other hand, at narrower stance, a higher control gain is needed (in both Bingham *et al* and our model). And, again, at sufficiently higher gain, their model becomes unstable. They find a smaller range of gains is stabilizing for wide stance as for narrow stance. Thus, they reason, that a control system that is challenged in its abilities to set gain levels, has an easier time setting the gain in narrow stance.

### Critical comment on objectivity in [3]

We do not know how a biological neural system quantifies gains. And, the concept of ‘narrower gain range’ depends on this measure. For example, the neural system might use the log of the engineering gain, or the reciprocal (the compliance). A range of gains that is smaller using engineering gain may not be smaller using these other measures. That is, the concept of ‘smaller gain range’ is not objective, in that it depends on how one chooses to measure gain (Unless one range totally encompasses the other, which is not the case in this model).

### Applicability of stability results to non-Parkinsons humans

In most (non-Parkinsonian) situations where balance is a challenge (neural ataxia), more-spread stances are chosen by both humans (wider) and horses (longer) [1, 4], not the narrower stance observed for standing human Parkinson’s patients. The Bingham *et. al.* [3] model, does not claim to be, and is not predictive in these other situations.

### Sway period in horses

The simplest rationale for our neglect of delay in our quantification of neural challenge is that the observed sway periods for standing horses are on the order of seconds, whereas the neural delays are probably on the order of tenths of second. For such motions, feedback with and without delay would have similar predictions. That is, in contrast with human grasp control, for postural control in horses the characteristic times of motions are slow enough so that neural delay may not be relevant.

This possible irrelevance of neural delay for standing horses is apparently not the case for standing humans, however, as witnessed by system identification [12] on humans being able to distinguish neural delay from instantaneous feedback.

## Appendix F: Parameters in the model

**Figure 6:**
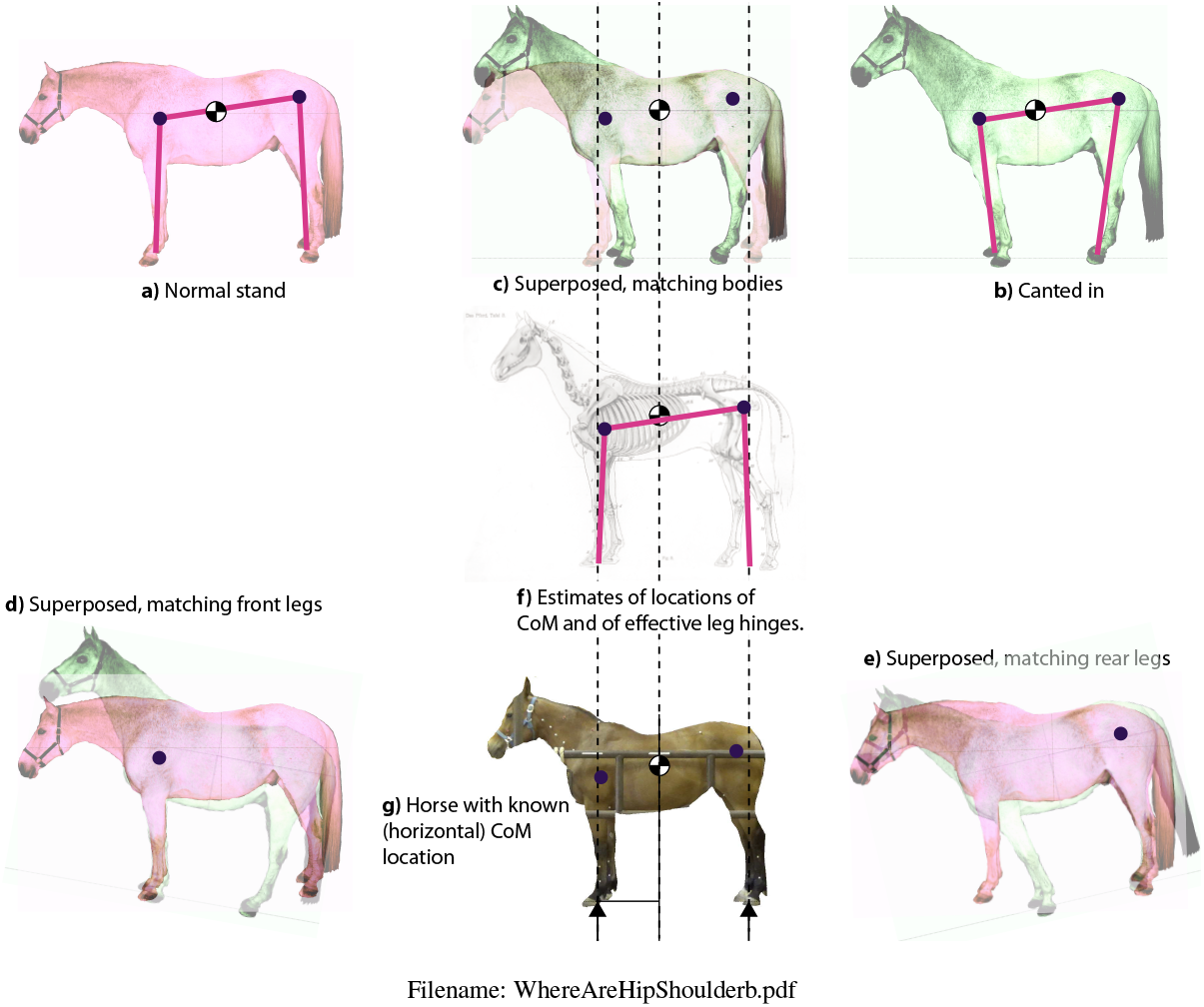
Locating the CoM and the effective shoulder and hip. One horse was photographed twice: **a)** once in normal posture and **b)** once with legs canted in. **c)** shown superposed. By rotating the photos **(d)** and**(e)**, locations are found for which relative rotation about those points best fits the observed motion of the leg relative to the body (d and e) and the body relative to the leg (c). **(g)** With a different horse, we found the center of mass using four force plates (unpublished data of Gellman K, Shoemaker JM, and Reese EM). All three horse images are rescaled to about the same size so that the effective hinges and the center of mass can be shown on all. For calculations, the dimensions for the effective leg lengths, back length, and center of mass location on the back, are measured from the photos, assuming a 16 hand (1 hand = 4”) horse. (photos by Judith M. Shoemaker)

For calculations we created a “standard horse” based on measurements from a few real horses. We scaled pictures to a common size so that pictures could be overlaid and compared. We scaled to a 16 hand horse, 64” from ground to withers (the high part of the back, just behind the neck).

### Location of shoulder and hip joints

Location of the effective shoulder joint and hip joint was done by superimposing photos of a single horse with more and less splayed legs. The effective shoulder is that point about which we can rotate one picture relative to the other so that in one case the legs are perfectly aligned and in the other the bodies are perfectly aligned. This point was found by sequential guessing using a graphics program (Adobe Illustrator). The hip was found similarly. All distances were measured from the scaled pictures.

### Parameter values

*m* = 500 kg. More or less typical for a 16 hand horse;
*g* = 10 m/s. Close enough to the standard value of 9.8 m/s^2^;
*ℓ_f_* = 1.05 m. Fore-leg length, ground to shoulder hinge;
*ℓ_r_* = 1.20 m. Rear leg length, ground to hip hinge;
*ℓ_b_* = 1.16 m. Body length, shoulder hinges to hip hinge
*l_CoM_* = 0.50 m. Distance along the shoulder to hip line from the shoulder hinge to the Center of Mass (CoM).
*l_CoM_* = 0.0 m. We assume the CoM was on the shoulder to hip line, thus the height of the CoM above that line is zero.

Leg splay *ℓ_g_* (also called *d*) is measured as the distance between the ground placement of the front and rear hooves. This parameter was varied.

## Appendix G: Collapsed Postures

This last example of canted-in posture is more tentative. Totally exhausted or traumatized humans can take on a characteristically collapsed, hunched-over posture. We see this posture in photos of prisoners of war, refugees fleeing a natural disaster or combat, and survivors of a traumatic event. Similarly, there is some indication that exhausted horses may have a characteristic canted-in posture. This is depicted in the sculptures “The Tired Hunter” by J. Willis Good (1875) and also the “End of the Trail”, by James Earle Fraser (1905) [8]. Both portray horses with narrow (canted-in) stance, and the latter also shows a rider with collapsed upper body posture. One possible explanation for these collapsed postures might be that conscious cognitive signals (intent) for upright posture are diminished in the exhausted individual, with a corresponding loss of flexor inhibition and extensor tone (Dr. Carl DeStefano — personal communication). Another possibility, (which may also apply to EMND) is that in these collapsed postures, the balancing torques are carried by bones and ligaments instead of muscles. In this explanation, these collapsed postures would not be chosen in normal circumstances because they are likely uncomfortable, possibly damaging to tissue, and not conducive to versatile functioning. One can view these mechanisms as failures of motor output.

